# Multiplexed phosphoproteomics of low cell numbers using SPARCE

**DOI:** 10.1101/2024.10.18.619068

**Authors:** Emily J. Gaizley, Xiuyuan Chen, Amandeep Bhamra, Silvia Surinova

## Abstract

Understanding cellular diversity and disease mechanisms requires global analysis of proteins and their modifications. While next-generation sequencing has advanced our understanding of cellular heterogeneity, it fails to capture downstream signalling networks. Ultrasensitive mass spectrometry-based proteomics enables unbiased protein-level analysis of low cell numbers, down to single cells. However, phosphoproteomics remains limited to high-input samples due to sample losses and poor reaction efficiencies associated with processing low cell numbers. Isobaric stable isotope labelling is a promising approach for reproducible and accurate quantification of low abundant phosphopeptides. Here, we introduce SPARCE (<u>S</u>treamlined <u>P</u>hosphoproteomic <u>A</u>nalysis of <u>R</u>are <u>CE</u>lls) for multiplexed phosphoproteomic analysis of low cell numbers. SPARCE integrates cell isolation, water-based lysis, on-tip TMT labelling, and phosphopeptide enrichment. SPARCE outperforms traditional methods by enhancing labelling efficiency and phosphoproteome coverage. To demonstrate the utility of SPARCE, we analysed four patient-derived glioblastoma stem cell lines, reliably quantifying phosphosite changes from 1,000 FACS-sorted cells. This workflow expands the possibilities for signalling analysis of rare cell populations.

## Introduction

Cells exhibit diverse phenotypes, behaviours and functions, governed by the proteome^1^. Analysing distinct cell populations is essential for unravelling disease mechanisms which arise from cellular dysfunction^2,3^. While next-generation sequencing has transformed the study of cellular heterogeneity, it cannot provide insights into the signalling networks responsible for cell decision-making^4^. Liquid chromatography coupled to tandem mass spectrometry (LC-MS/MS) is a powerful technique for the unbiased quantification of proteomes and their proteoforms^5^. However, the analysis of phosphorylated proteins, which requires an input-sensitive phosphopeptide enrichment step, has historically been limited to the study of high microgram to low milligram protein amounts due to significant challenges associated with processing limited material^6^.

Given the extensive sample loss associated with multistep processing of nanogram scale samples, there have been limited attempts to integrate phosphoproteomics with existing cell isolation techniques, which are required to obtain pure cell populations of biological importance^7^. Fluorescence-Activated Cell Sorting (FACS) is a single-cell isolation technique widely used to analyse and sort distinct cell populations based on cell-specific fluorescent markers and morphological parameters. Its precision and versatility make it an important tool for both basic research and clinical applications^8^. Despite its utility, the high volumes and buffer salinity associated with sorted populations are generally incompatible with current lysis and isobaric labelling methods. Traditional protein extraction workflows favour chaotropic or detergent-based buffers followed by sonication, reduction and alkylation, dilution and pH adjustment^9^. This approach, originally designed for high-input bulk proteomics of millions of cells, has been adapted by Amon *et al*., for the proteomic analysis of 25,000 human hematopoietic FACS-sorted cells^10^. In this study, they highlighted the need for careful reduction of volumes after sorting and its critical impact on reproducibility. At the opposite end of the scale, single-cell proteomic workflows have employed water-based buffers to achieve hypotonic lysis followed by a freeze-heat cycle to denature proteins^11,12^. Given that nanogram-range phosphoproteomics faces similar challenges to single-cell proteomics, water-based lysis might yield deeper proteome coverage compared to traditional urea-based methods for low cell numbers.

To obtain reliable phosphopeptide quantification from a few thousand cells, multiplexed isobaric stable isotope labelling is emerging as one of the most effective approaches offering more accurate and reproducible quantification of low abundant peptides compared to label-free techniques^13^. Multiplexing presents the opportunity to include a carrier proteome sample which typically resembles or is a mixture of the individual samples. The carrier proteome channel ‘boosts’ the MS1 signal, thereby triggering more MS2 scans and identifying more peptides^14,15^. Using this approach, Tsai *et al*., developed a tandem tip-based method combining C18-IMAC-C18 desalting and phosphopeptide enrichment of TMTpro-labelled peptides, which enabled the quantification of ∼600 phosphopeptides from 100 sorted cells^7^. This work demonstrates that multiplexed phosphoproteomic analysis of small numbers of FACS-sorted cells is possible.

While multiplexing is a promising approach for low cell number phosphoproteomics, standard TMT labelling protocols typically require a hundred of micrograms of input material and are generally performed in-solution by adding the TMT reagent to concentrated, purified peptides^16^. This sample preparation format is incompatible with low-input samples due to the low substrate concentration, which results in reduced labelling efficiency. This can be partially overcome by reducing the reaction volume and increasing the amount of TMT, but this can result in excess TMT reagent and overlabelling, which can inhibit peptide ionisation and increase sample complexity^17,18^. Performing TMT labelling on-tip instead of in-solution enhances reaction kinetics and reduces sample loss of low input samples^18-20^. On-tip labelling has the potential to streamline low-input sample preparation workflows by combining labelling with desalting to limit sample loss and concentrate peptides. A handful of studies have already demonstrated the success of this method, with some highlighting its application for low-input phosphoproteomics. On-tip TMT labelling was first demonstrated using C18 cartridges for high-input samples but has since been scaled down for low-input amounts using C18 tips^18,19,21^. Park *et al*. recently published their one-STAGE tip method for TMT labelling of five cells^20^. This approach integrates protein extraction and digestion with on-tip TMT labelling, eliminating the need for sample transfer. While this method has not yet been combined with phosphopeptide enrichment, it demonstrates the potential of on-tip TMT labelling for processing low cell numbers.

Here, we present SPARCE (<u>S</u>treamlined <u>P</u>hosphoproteomic <u>A</u>nalysis of <u>R</u>are <u>CE</u>lls), an end-to-end multiplexed phosphoproteomic workflow for the analysis of FACS-sorted cells (**Fig. 1**). SPARCE addresses the challenges associated with low-input phosphoproteomics by simplifying and integrating: (i) water-based lysis of FACS-sorted cells, (ii) one-step protein digestion, (iii) on-tip desalting and TMTpro labelling, with (iv) phosphopeptide enrichment of multiplexed samples. SPARCE significantly enhances the proportion of fully labelled peptides while maintaining high labelling efficiency and improved peptide identifications. We demonstrate that SPARCE outperforms traditional lysis and labelling techniques and yields improved phosphopeptide identifications as a result of improved reaction efficiency, reduced sample loss, and better retention of hydrophilic peptides. Finally, we validated the quantitative accuracy of SPARCE in patient-derived glioblastoma stem cell lines. By comparing results from 1,000 cells and 20,000 cells, we show that SPARCE is capable of quantifying reproducible phosphopeptide changes across cell lines to yield biologically meaningful results. This workflow expands the possibilities for phosphoproteomic analysis of rare cells, making it a valuable tool for researchers looking to combine FACS with multiplexed LC-MS/MS.

**Fig. 1.**
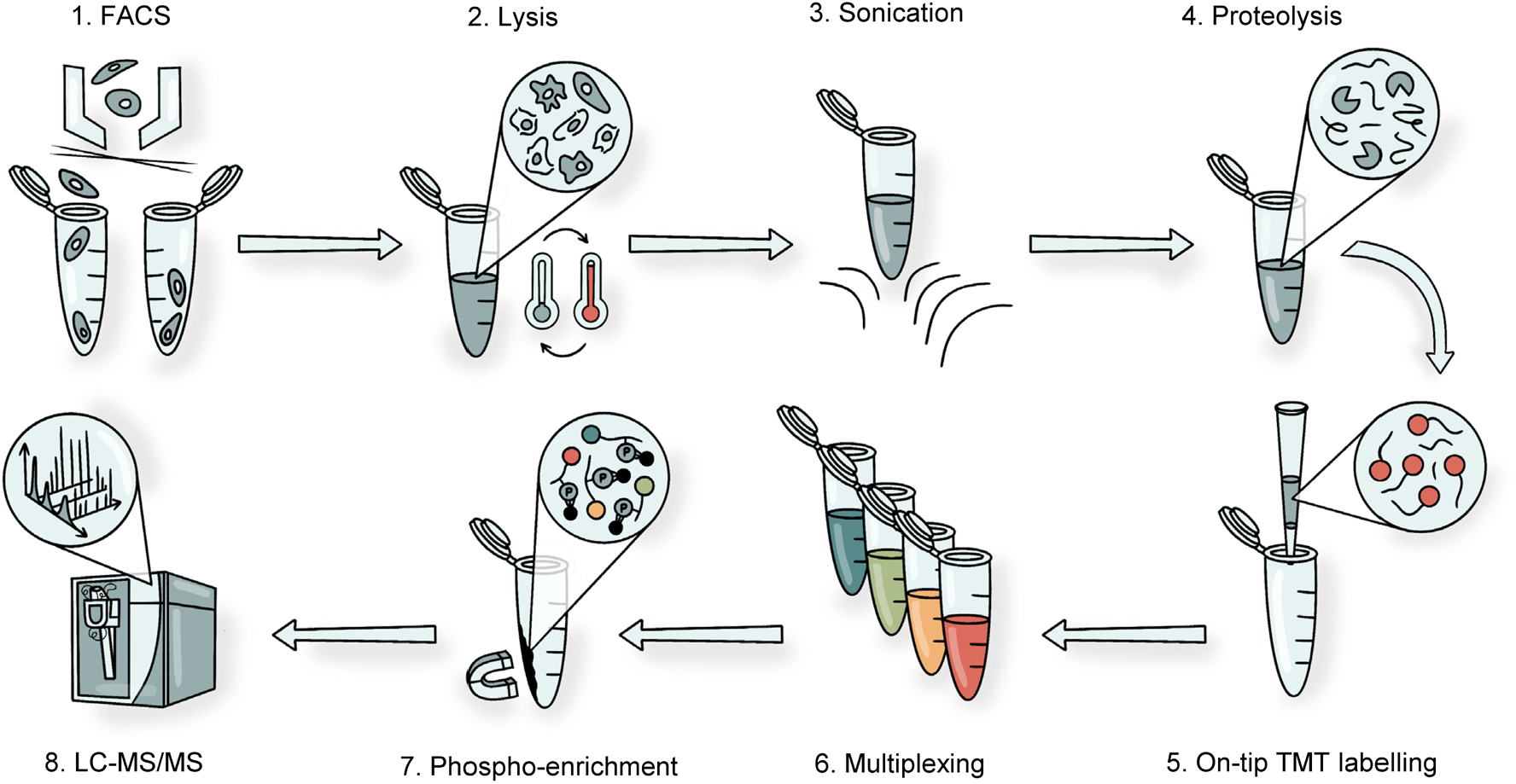
Schematic of the SPARCE workflow. Cells are isolated using FACS and subjected to lysis by freeze-heat cycles and sonication. Proteins are digested, and peptides are labelled and desalted on the same tip. Labelled samples are multiplexed, and phosphopeptides are enriched for LC-MS/MS analysis.

## Results

### FACS-compatible water-based lysis outperforms traditional methods

During cell sorting by FACS, cells are suspended in sheath fluid, typically phosphate-buffered saline (PBS), to maintain the osmotic pressure and preserve cell viability. The volume of PBS collected after cell sorting represents a major challenge for low-input sample preparation. Attempting to remove the buffer necessitates careful optimisation to prevent excess sample loss and ensure reproducibility^10^. Hence, we opted to isolate and sort live single cells (**Supplementary Fig. 1a**) directly into a lysis buffer.

To mitigate the need for buffer exchange or dilution prior to digestion, 1,000 glioblastoma stem cells, corresponding to approximately 75 ng protein (**Supplementary Fig. 1b**), were FACS-sorted directly into 20 μL water to achieve hypotonic lysis. Immediately after sorting, cells were stored on dry ice to limit further enzymatic modifications. The cells were lysed with three freezing and heating cycles; each cycle included five minutes on dry ice and five minutes at 90°C. Following lysis, samples were cooled down to room temperature and sonicated for 15 minutes to ensure efficient cell lysis. Using this method, the number of fully digested peptides increased fivefold when compared to a typical urea-based lysis protocol (**Fig. 2a**). In this case, 200 ng of trypsin at a ∼2:1 protease-to-protein ratio was used to ensure trypsin was in excess. The increase in identifications is most likely due to reduced unspecific sample loss and improved digestion efficiency, given that the total volume of the water-based sample remains largely unchanged compared to the urea-based sample that requires dilution prior to digestion. Next, we evaluated the use of protease and phosphatase inhibitors that are traditionally used in phosphoproteomic workflows. Their inclusion significantly impeded digestion, resulted in large amounts of undigested protein, affected the overall system performance, and led to a three and half fold drop in identified peptides (**Supplementary Fig. 2a**). Based on these findings, inhibitors were omitted from subsequent experiments.

**Fig. 2.**
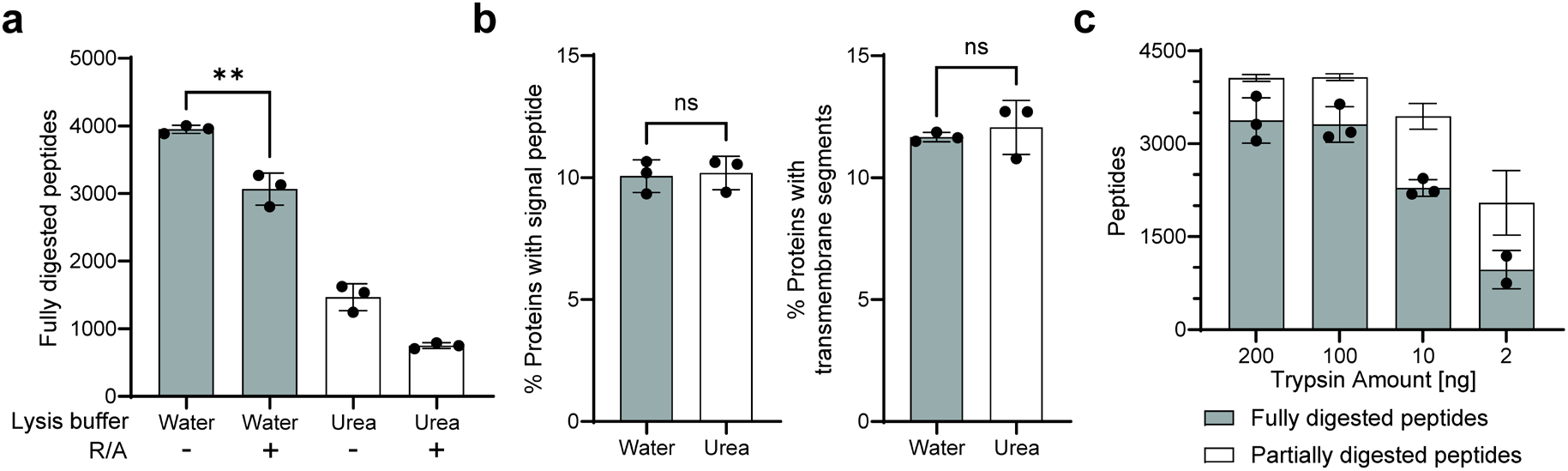
Optimisation of FACS-compatible lysis for low-input samples. **a** The effect of different lysis methods on the digestion efficiency of 1,000 cells using 200 ng trypsin (2:1 protease-to-protein ratio). R/A, reduction and alkylation. **b** Assessment of the possible detection bias in proteins with signal peptides (left), or transmembrane segments (right) for different lysis buffer methods. **c** Optimisation of trypsin amount for the digestion of 1,000 cells.

The breaking of disulfide bonds by reduction and alkylation of cysteine residues has been a mainstay in traditional proteomic sample preparation workflows^22^. The omission of reduction and alkylation prior to trypsin digestion increases the complexity of MS2 spectra when two peptides are linked by disulfide bonds, thus limiting the identification of cystine-containing peptides. Until recently, this step has broadly been accepted as a prerequisite for the processing of proteomic samples. However, we observed a significant decrease (∼25%) in fully digested peptides following reduction and alkylation, suggesting that trypsin efficiency may be impaired in the presence of reduction and alkylation agents (**Fig. 2a**). We speculate that this is caused by the structural destabilisation of trypsin following the reduction of its disulphide bonds by TCEP. Hence, given the deleterious effect of TCEP on digestion efficiency of nanogram scale input samples, this step was omitted from future experiments.

To determine whether we could extract membrane-bound proteins using the water-based lysis method in the absence of detergents or chaotropic agents, we analysed the distribution of proteins identified across the two methods. There was no significant difference in the percentage of extracellular and transmembrane proteins detected in the urea-based method and the water-based method (**Fig. 2b**). Hence, the water-based lysis method can sufficiently extract non-cytosolic proteins, which is important for capturing the full diversity of proteins in discovery-based proteomic experiments. While the ratio of transmembrane and extracellular proteins identified did not change significantly, the absolute number increased by around 30% in the water-based method, indicating this is indeed the superior approach for low-input samples (**Supplementary Fig. 2b**).

Following lysis buffer optimisation, we sought to determine the optimal concentration of trypsin to achieve complete digestion. Typical protease-to-protein ratios range from 1:10 to 1:100, however, this may not be sufficient to overcome the challenge of poor substrate concentrations in low-input samples. Indeed, increasing the protease-to-protein ratio to 50:1 has been shown to significantly improve peptide identifications in single-cell proteomic studies^23^. Here, we observed increased digestion efficiency and enhanced reproducibility at ratios as high as ∼2:1, with improvements plateauing at a ratio of ∼1:1. Compared with 2 ng trypsin (∼1:50, protease-to-protein ratio), which is commonly used in conventional proteomics, 100 ng trypsin doubled both the number of peptide identifications as well as the percentage of fully digested peptides (**Fig. 2c**). Based on these results, we opted to use 100 ng trypsin for subsequent experiments.

### Combining desalting and labelling on-tip leads to superior labelling efficiency and the retention of hydrophilic peptides

The high salinity of the sheath fluid used during FACS means that desalting is required following proteolysis. On-tip TMT labelling is an attractive alternative to in-solution methods because it enables desalting in conjunction with labelling, thus streamlining the workflow and reducing sample losses. On-tip labelling has been previously performed on phosphopeptides enriched from 1-100 μg of HeLa digests using TMT10plex reagents^19^. Here, we sought to establish on-tip labelling for nanogram-scale samples with TMTpro reagents.

It is important to optimise TMT labelling to achieve high labelling rates on α- and ε-amino groups on peptide N termini and lysine side chains and maintain low partial labelling rates to minimise an increase in analyte complexity. Equally, it is important to minimise overlabelling on the hydroxyl group on serines, tyrosines and threonines, as well as on the imidazole group on histidines. Another reason for careful optimisation of the required TMTpro reagent was to reduce the levels of unreacted and/or hydrolysed reagents as much as possible to prevent adverse effects on column binding capacity and peptide ionisation. Using 1,000 cells, we labelled the samples with different TMTpro amounts, ranging from 0.3 to 12 μg. While 0.3 μg yielded the most peptide identifications, the rate of labelling was low, therefore, 3 μg was selected for subsequent experiments due to improved reproducibility and low rate of overlabelling (**Fig. 3a, b**). We further tested 3 μg of reagent on different input sample amounts ranging from 500 to 40,000 cells (equivalent to approximately 37.5 ng to 3 μg) and demonstrated stable labelling efficiencies, partially labelling rates and over labelling rates over the range (**Fig. 3c**).

**Fig. 3.**
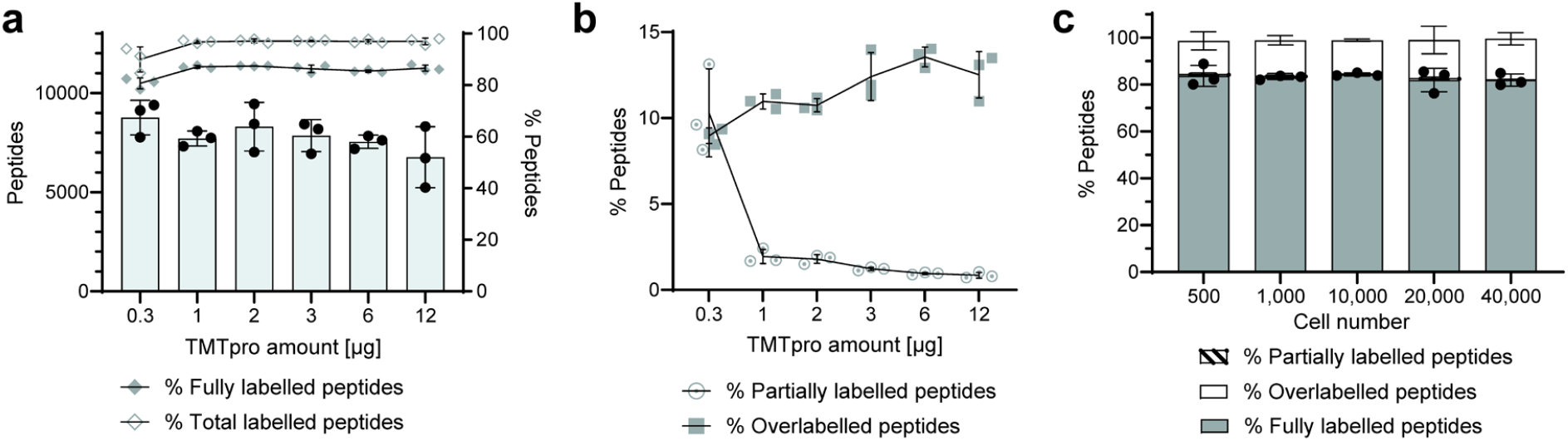
On-tip labelling method optimisation for low-input samples. **a** Peptide identifications and labelling efficiency for varying amounts of TMTpro. **b** The overlabelling and partially labelling rates of different TMTpro amounts during labelling. **c** On-tip labelling of different cell numbers using 3 μg TMTpro. Fully labelled peptides indicate peptides where all available residues have been labelled, partially labelled peptides are the peptides with some available sites for labelling, and overlabelled peptides are the peptides with off-target labelling.

Following optimisation of the on-tip labelling method, we compared it with in-solution labelling. While we used 3 μg of TMTpro reagent for on-tip labelling, 24 μg of TMTpro was additionally used for in-solution labelling to ensure the TMT amount was not limiting. Our optimised on-tip method significantly reduced sample loss, almost doubling the number of peptide identifications compared to the in-solution method (**Fig. 4a**). The on-tip method demonstrated superior labelling efficiency and a low percentage of partially labelled peptides (**Fig. 4b**). Moreover, the on-tip method retained significantly more hydrophilic peptides since the percentage of hydrophilic peptides increased by ∼5% (**Fig. 4c, Supplementary Fig. 3**). This is of particular importance in the context of downstream phosphopeptide enrichment and is likely attributable to the labelling being performed before the desalting wash step. This is evidenced in **Fig. 4d**, where hydrophilic peptides account for 75.9% of the peptides uniquely identified with the on-tip method.

**Fig. 4.**
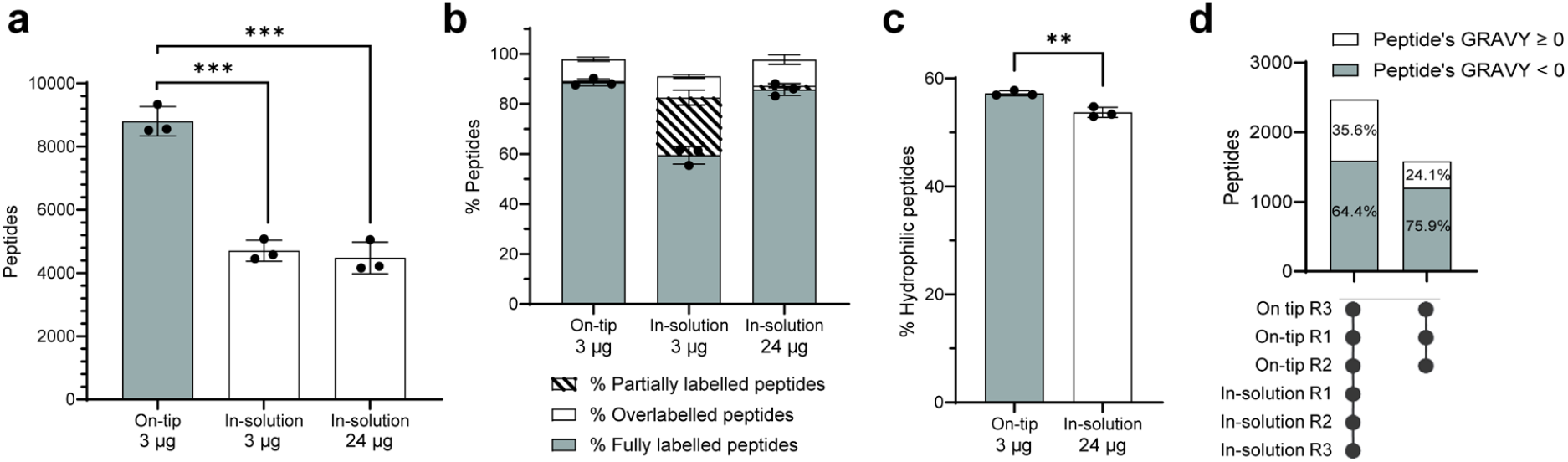
Comparing on-tip and in-solution labelling methods. **a** Identified peptides in on-tip versus in-solution method. **b** The labelling output between the on-tip and in-solution method. **c** The percentage of hydrophilic peptides in the peptide groups detected by the two methods. The peptides with a GRAVY factor less than 0 represent hydrophilic peptides. **d** Hydrophobic distribution of specific peptide groups. The first column shows the distribution of peptide detected in all six samples, and the second column shows the distribution of peptide exclusively detected in all three samples using the on-tip labelling method. 24 μg TMTpro was used for in-solution labelling.

### Low-input multiplexed phosphoproteomic analysis of patient-derived glioblastoma cell lines

It is important that new methods can be directly applied to relevant biological samples and achieve robust coverage of their phosphoproteomes. To test the quantitative accuracy and reproducibility of SPARCE in a relevant model system, we analysed four patient-derived glioblastoma stem cell lines, each representing a distinct molecular subtype^24,25^. For each cell line, we FACS-sorted 1,000 and 20,000 cells in triplicate (herein referred to as low and high plex, respectively) (**Fig. 5a**). By varying the multiplexed amount of input material, we determined that 160,000 cells were required to identify >1,800 phosphorylated peptides (**Supplementary Fig. 4**). To ensure the multiplexed cell numbers were similar across the high and low plexes, a carrier channel of 160,000 sorted cells representing a mixture of all the biological samples was included with the individual 1,000 cell samples (**Supplementary Fig. 5a**). To improve the MS2 spectral quality, the MS2 mass range was adjusted to start acquisition at 126.5 *m/z* to omit the carrier channel in the 126 position, as previously described^26^. This analysis yielded > 2,500 and > 1,500 quantified phosphosites per condition for the high and low plexes, respectively (**Fig. 5b**).

**Fig. 5.**
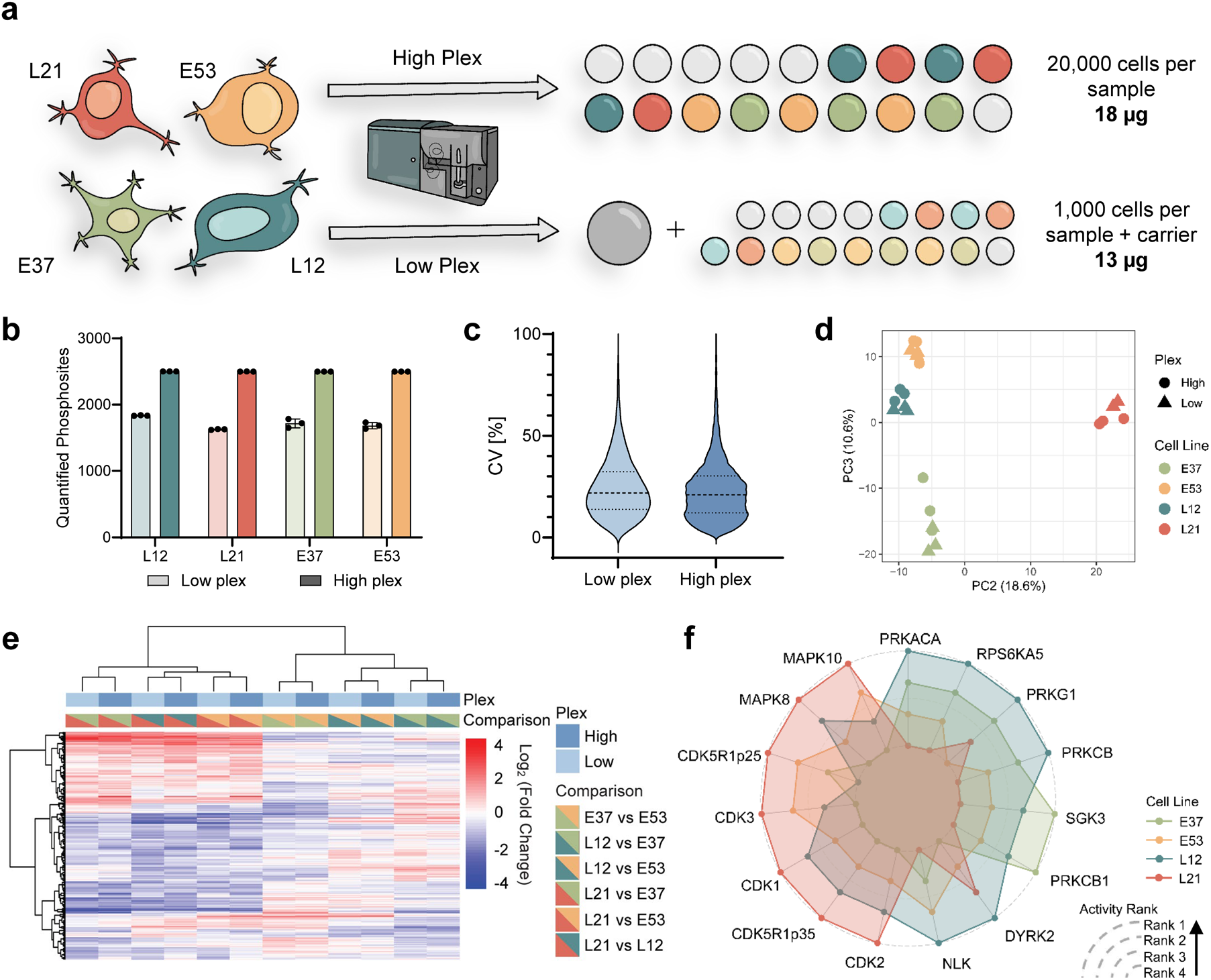
Comparison of the SPARCE workflow using different input amounts. **a** Experimental design to investigate the differences between four patient-derived glioblastoma cell lines. The plex with a carrier was named as “low” plex, while the “high” plex did not use a carrier. **b** The number of phosphosites quantified from each cell line in both plexes. **c** The distribution of CV values for quantified phosphosites from both plexes. The datapoints with a CV value between 0% to 100% were shown. The dotted lines present the first and third quartiles, while the dashed lines are the position of the median value. **d** The PCA results based on phosphosite abundance, x-axis is PC2 and the y-axis is PC3. **e** The heatmap of differential phosphosites between different cell lines found from both plexes, with column-based clustering. **f** The kinase activity profiles of different cell lines, inferred and ranked using PTM-SEA.

To examine the reproducibility of SPARCE, coefficients of variation (CVs) were calculated for each quantified phosphopeptide in each cell line, and the results were plotted for both the low and high plex experiments. The median CV values for the low and high plexes were 21.8% and 20.9%, respectively (**Fig. 5c, Supplementary Fig. 5b**), demonstrating comparable precision of quantification across the two plexes. We further assessed phosphosite abundances across high and low plexes for each of the cell lines. We observed clear clustering of the cell lines across the plexes, indicating high reproducibility and quantitative precision (**Fig. 5d, Supplementary Fig. 5d**). By carrying out pairwise differential abundance testing across the cell lines, we aimed to understand whether phosphosite regulations were also reproducible between the low and high plex experiments. Unsupervised hierarchical clustering revealed reproducible differentially abundant phosphosites, with high and low plexes of the same condition comparisons clustering together (**Fig. 5e**). This was further validated by calculating Spearman correlation coefficients, demonstrating excellent correlations across plexes for the same comparisons (**Supplementary Fig. 5c**).

Next, we aimed to evaluate the quality of functional insights that could be derived from SPARCE, and to determine the extent to which patient-derived cell lines exhibit differential kinase activities. We used PTM-SEA^27^ with the iKiP database to infer kinase activities from differentially abundant phosphosites across cell line comparisons in both plexes (**Supplementary Fig. 5e**). Using these results, we sought to determine the extent to which patient-derived cell lines exhibit differential kinase activities by ranking the cell lines in order of activity for each kinase (**Fig. 5f**). This analysis revealed that cell line L21 has the highest activity of MAPK10, MAPK8, p25, p35, CDK1, CDK2 and CDK3. These kinases are involved in cell cycle regulation and stress response mechanisms. On the other hand, cell line L12 and E37 exhibit increased activity of kinases associated with signalling pathways that regulate cellular responses to external stimuli (such as hormones and growth factors) and intracellular second messengers (such as cAMP, cGMP, and PI3K). These pathways are heavily involved in metabolism, cell survival, and growth regulation. This relative comparison of global kinase activities across patient-derived cell lines highlights that it is indeed possible to gain meaningful insights into the phosphoproteome with limited material. This has implications for understanding patient-specific phenotypes associated with distinct signalling programmes and expands the possibility for more personalised approaches to treatment. Taken together, our data demonstrate that SPARCE is a powerful method for probing global phosphoproteome changes in low-input samples.

## Discussion

Here, we present SPARCE, a method for streamlined phosphoproteomic analysis of low cell numbers. Importantly, SPARCE is compatible with FACS, which is widely used for the isolation of subpopulations of cells. Lysis buffer optimisation experiments revealed that a water-based buffer in conjunction with freeze-heat cycles outperforms traditional urea-based lysis methods. This is likely due to the high volumes associated with dilution of urea prior to digestion. In addition, we found that removing protease inhibitors, and reduction and alkylation agents further improved peptide identifications due to the negative effects on trypsin activity. Increasing trypsin concentrations further improves the number of peptide identifications, highlighting the importance of optimising digestion for low-input samples where low enzyme-substrate concentration is the limiting factor. Together, these findings highlight the importance of low volumes and minimal processing steps for effective digestion and improved recovery of low-input samples.

SPARCE addresses the limitations of traditional phosphoproteomic workflows by nearly doubling peptide identifications compared to in-solution methods. SPARCE effectively integrates TMT labelling and desalting, minimising sample loss and increasing labelling efficiency. The on-tip method significantly retains more hydrophilic peptides, which is important for downstream phosphopeptide enrichment, given the hydrophilic nature of phosphopeptides.

Using SPARCE, we demonstrated the utility of the method by performing multiplexed phosphoproteomic analysis on four patient-derived GBM cell lines. Multiplexing is a major advantage of TMT labelling and the use of ‘carrier’ channels has shown promise in the field of single-cell proteomics, where throughput and protein amount are key limitations^11,28,29^. However, this has not been without criticism due to the reduced quantitative accuracy and increased ratio compression associated with carrier channels. Acceptable carrier ratios have been reported ranging from <20x to <200x depending on the instrument parameters, type of sample and number of cells in the sample of interest^26,30-32^. Hence, we opted to use a 160x cell carrier channel for the low plex experiment. CV calculations revealed comparable values (approximately 20%) for both the low and high plex experiments, indicating that the carrier channel does not negatively impact the quantitative accuracy in this context.

Overall, the ability of SPARCE to reproducibly quantify phosphopeptide changes, from as few as 1,000 cells per sample, makes it a valuable tool for researchers and expands the possibilities for routine phosphoproteomic analysis of rare cells without the need for additional specialised equipment. By combining FACS with LC-MS/MS, SPARCE offers a new frontier in uncovering cellular functions and signalling pathways that are critical to understanding complex biological systems.

## Methods

### Cell culture

The patient-derived GSC lines (GCGR-E37, GCGR-E53, GCGR-L12 and GCGR-L21) were cultured and maintained in serum-free GSC media as an adherent monolayer, as previously described^33^. Briefly, the cells were cultured in DMEM/HAMS-F12 (Sigma D8437), supplemented with 1.45% (v/v) glucose (Sigma, USA), 1% (v/v) 100× MEM Non-Essential Amino Acids Solution (Gibco, USA), 1% (v/v) pen-strep (Gibco, USA), 0.16% (v/v) BSA soln 7.5% (Gibco, USA), 0.2% (v/v) 50 mM 2-mercaptoethanol(Gibco, USA), 1% (v/v) 50× B27 supplement (Gibco, USA), 0.5% (v/v) 100× N2 supplement (Gibco, USA), 10 ng/mL (final concentration) mouse epidermal growth factor (PeproTech, USA), 10 ng/mL (final concentration) human fibroblast growth factor-basic (PeproTech, USA), and 2 μg/mL (added immediately prior to use, final concentration) laminin (Sigma, USA). Flasks (Fisher Scientific, USA) were pre-coated with 5 μg/mL laminin in PBS (Gibco, USA) for 1 hour before plating the cells. Media were changed every 5 days, and cells were split at 80-90% confluency. Cells were dissociated using Accutase solution (Sigma, USA), centrifuged and washed before re-plating. GCGR-E37 was used for all technical optimisation experiments, and GCGR-L12, GCGR-L21, GCGR-E37, and GCGR-E53 were used for the cell line comparison experiment, herein referred to as L12, L21, E37 and E53, respectively.

### Cell sorting and lysis

BD FACS Aria Fusion/III (BD Biosciences, USA) was used to sort live cells with a 100 μm nozzle. Before sorting, cells were washed twice with PBS and dissociated using the Accutase solution (Fisher Scientific, USA). Cells were centrifuged, washed twice with PBS and resuspended in Hoechst 33258 (Invitrogen, USA, 1:10000 with PBS). Cells were sorted into lysis buffer, which was added to 1.5 mL Protein LoBind Tubes (Eppendorf, UK).

For the water-based lysis method, cells were sorted directly into 20 μL LC-MS grade water (Fisher Scientific, USA). The cells were lysed after being subjected to three freezing and heating cycles. During each cycle, cells were snap frozen using dry ice for five minutes followed by immediate heating at 90°C in a thermomixer (Eppendorf, Germany) without shaking for five minutes. Following this, samples were cooled down to room temperature and sonicated for 15 min at 150 W using an ultrasonic water bath (Fisher Scientific, USA).

For the urea-based lysis method, cells were sorted directly into 20 μL 8 M urea (Sigma, USA) in 50 mM Triethylammonium bicarbonate (TEAB, Sigma, USA) and sonicated for 15 min at 150 W using an ultrasonic water bath (Fisher Scientific, USA).

Protease (Mini cOmplete™ EDTA-free Protease Inhibitor Cocktail, Roche, USA) and phosphatase (PhosphoSTOP, Roche, USA) inhibitors were used only where stated and were added to the lysis buffer according to the manufacturer’s recommended concentration and protocol. In summary, 1/5 tablet of Protease Inhibitor Cocktail and 1 tablet of PhosphoSTOP were added into 10 mL lysis buffer, for both water-based and urea-based lysis buffers.

### Micro BCA assay

1,000, 5,000, 10,000 and 20,000 cells were sorted in triplicate, lysed using the water-based lysis method and dried using a vacuum concentrator (Labconco, USA). Samples were resuspended in 40 μL LC-MS grade water, and the micro BCA protein assay working buffer was added according to the kit manufacturer’s protocol (Thermo Scientific, USA). Absorbance was measured for samples and standards at 562 nm using a NanoDrop 2000 (Thermo Scientific, USA). The standard curve was calculated in GraphPad Prism version 10.1.2 for Windows (GraphPad Software, USA).

### Protein digestion

Protein reduction and alkylation reagents were only used where mentioned. For reduction, samples were treated with 100 mM Tris(2-carboxyethyl)phosphine hydrochloride (TCEP, Sigma, USA) dissolved in 50 mM TEAB buffer, achieving a final TCEP concentration of 5 mM. This was followed by an incubation period of 20 min at 37°C. For alkylation, 200 mM 2-Chloroacetamide (CAM, Sigma, USA) prepared in 50 mM TEAB buffer was added, resulting in a final CAM concentration of 5 mM. The samples were then incubated for 20 min at room temperature in the dark.

For samples processed using the urea-based lysis method, 50 mM TEAB was used to dilute the final urea concentration to <1.5 M before digestion.

Trypsin gold (Promega, USA) was resuspended with 100 mM TEAB to a 40 ng/μL concentration. Unless stated otherwise, for samples containing 10,000 or fewer cells, a protease-to-protein ratio of 1:1 was used, and for cell numbers higher than 10,000, 1:10 was used. Samples were incubated with trypsin at 37°C for 4 hours at 500 rpm and thereafter at room temperature overnight without shaking. 1% trifluoroacetic acid (TFA, Fisher Scientific, USA) was added to stop the reaction (0.1% final concentration) and adjust the pH for subsequent loading onto ZipTip columns (Sigma, USA) for either labelling or desalting.

### In-solution TMTpro labelling

Samples were desalted prior to in-solution labelling using ZipTip C18 columns with a 0.6 μL bed (Sigma, USA) by means of a centrifuge adaptor (GL Science, USA). Firstly, 50 μL LC-MS grade acetonitrile (ACN, Fisher Scientific, USA) was passed through the ZipTip columns using 110 g centrifugation to activate the C18 material. The columns were then washed once with 50 μL 0.1% TFA in water to equilibrate the column. To maximise peptide binding, samples were loaded three times by passing the flow through back on to the column. The columns were then washed with 50 μL 5% ACN/0.1% TFA in LC-MS grade water. Peptides were eluted twice with 50 μL 50% ACN/0.1% TFA in water. Samples were lyophilised using a vacuum concentrator (Labconco, USA).

Peptides were resuspended with 10 μL LC-MS grade water by vortexing at 17,000 *g* for 30 min, followed by sonication for 15 min in the ultrasonic water bath. Samples were centrifuged again at 17,000 *g* for 5 min. Anhydrous acetonitrile (Sigma, USA) was used to resuspend the TMTpro zero reagent (Thermo Scientific, USA) at 3 μg/μL. The required amount of TMTpro zero, either 3 or 24 μg, was added to the sample keeping the reagent volume consistent at 8 μL. Samples were labelled for 1 h at room temperature. The excess TMT reagent was quenched using 0.04% hydroxylamine (final concentration, Sigma, USA) and incubated for 15 min at room temperature. A second desalting process was carried out as above.

### On-tip TMTpro labelling

The on-tip TMTpro zero and TMTpro labelling method combined desalting and labelling to streamline the workflow and reduce sample losses. The steps up to, and including, sample loading were the same as described in the section “In-solution TMTpro labelling”. After sample loading, phosphate buffer was prepared by diluting 1 M potassium phosphate monobasic solution (Sigma, USA) to 50 mM with LC-MS grade water and adjusting pH to 6.5 with 1 M TEAB. Next, phosphate buffer was used to wash the column to adjust the pH to nearly neutral. TMTpro zero or TMTpro was resuspended in the phosphate buffer to the desired concentration to ensure that 10 μL was consistently loaded onto the tip. The tip was transferred to a new tube, and 10 μL TMTpro zero or TMTpro reagent was loaded onto the tip and centrifuged at 110 g. The amount of TMTpro reagent used varied from 0.3 to 12 μg depending on the experiment.

During the one-hour labelling reaction, the tip was kept wet in phosphate buffer by reloading the flowthrough and ensuring the liquid was visible at both sides of the C18 bed. After the reaction, 40 μL 0.1% TFA was used to acidify the sample, and the labelled sample was re-loaded on the tip twice to recover any peptides from the flow through. The column was washed with 5% ACN/0.1% TFA to remove salts and other hydrophilic contaminants, and the elution step was performed twice. Samples were lyophilised using a vacuum concentrator (Labconco, USA).

### Phosphopeptide enrichment

Labelled samples were resuspended in the loading buffer (0,1M glycolic acid in 80% ACN, 5% TFA) and vortexed for 30 min prior to multiplexing. The loading buffer volume is variable depending on the number of samples, and the final volume of multiplexed samples was kept the same (200 μL). Phosphopeptide enrichment was performed using 10 μL (200 μg) Zr-IMAC HP beads (Resyn Biosciences, South Africa), according to the manufacturer’s instructions, with the only exception that TFA was used instead of FA to acidify the sample after elution. Post-enrichment desalting was carried out using BioPureSPN Mini, PROTO 300 C18 spin columns (Nest Group, USA). 600 μL LC-MS grade acetonitrile (ACN, Fisher Scientific, USA) was used to activate the C18 material, and then 600 μL 0.1% TFA in LC-MS grade water was used to equilibrate the columns. Samples were loaded three times to maximise peptide binding. The columns were washed with 600 μL 5% ACN/0.1% TFA. Peptides were eluted with 75 μL 50% ACN/0.1% TFA three times and lyophilised using a vacuum concentrator (Labconco, USA).

### Liquid Chromatography-Mass Spectrometry

All label-free experiments evaluating lysis and digestion conditions were performed on a Q-Exactive Plus mass spectrometer interfaced with a Nanospray Flex ion source and coupled to Easy-nLC 1200 (Thermo Scientific, USA). Peptides were separated on a 24 cm fused silica emitter, 75 μm ID, packed in-house with Reprosil-Pur 200 C18-AQ, 2.4 μm resin (Dr. Maisch, Germany) using a linear gradient from 5% to 30% acetonitrile/0.1% formic acid over 30 min at a flow rate of 250 nL/min. Precursor ions were measured in data-dependent mode at a resolution of 70,000 and AGC target value of 3 × 10^6^ ions. The ten most intense ions from each MS1 scan were isolated, fragmented in the higher-energy collisional dissociation cell, and measured in the orbitrap at a resolution of 17,500.

All experiments with TMTpro zero/TMTpro were analysed in DDA mode on an Orbitrap Exploris 480 mass spectrometer interfaced with a Nanospray Flex ion source and coupled to Easy-nLC 1200 (Thermo Scientific, USA). Peptides were separated using an Aurora Ultimate 25 cm × 75 μm C18 UHPLC column (IonOpticks, Australia), housed in a column oven (Sonation GmbH, Germany) set at 37 °C, with a two-stage gradient. The first stage involved increasing the acetonitrile/0.1% formic acid concentration from 8% to 21.6% over 50 minutes, followed by a second stage where the concentration was further increased from 21.6% to 32% over 10 min, at a flow rate of 150 nL/min.

For TMTpro zero samples, the MS1 scan was acquired at 120,000 resolution targeting ions in the scan range 375-1200 *m/z*. The Normalised AGC target was set to 300% (3e6 ions) (max. MS1 injection time set to “Auto”). The precursor ions were isolated with 0.7 *m/z* isolation window and fragmented with a Normalised Collision Energy (NCE) of 32. The MS2 scans were acquired at 45,000 resolution with the First mass set to 110 *m/z*. The Normalised AGC target set to 50% (5e4 ions) (max. MS2 injection time set to “Auto”) and the duty cycle time was set to 2 sec.

For TMTpro samples, the MS1 scan was acquired at 120,000 resolution targeting ions in the scan range 375-1200 *m/z*. The Normalised AGC target was set to 300% (3e6 ions) (max. MS1 injection time set to “Auto”). The Top12 precursor ions were isolated with 0.7 *m/z* isolation window and fragmented with a Normalised Collision Energy (NCE) of 32. The MS2 scans were acquired at 60,000 resolution with the First mass set to 126.5 *m/z*. The Normalised AGC target set to 90% (9e4 ions) (max. MS2 injection time set to 150 ms).

### Mass Spectrometry Data Analysis

Label free and TMTpro zero labelled data were searched using MaxQuant (v 2.3.1)^34^ against the human proteome sequence database retrieved from SwissProt on November 6, 2020. For LFQ data, Oxidation (M), Carbamidomethyl (C) and Acetylation (protein N-term) were set as variable modifications. For TMTpro zero data without phosphopeptide enrichment, TMTpro zero (K), TMTpro zero (peptide N-term), TMTpro zero (H, S, T), Oxidation (M) and Acetylation (protein N-term) were set as variable modifications. For TMTpro zero data with phosphopeptide enrichment, TMTpro zero (K), TMTpro zero (peptide N-term), TMTpro zero (H), Phosphorylation (S, T, Y), Oxidation (M) and Acetylation (protein N-term) were set as variable modifications. Enzyme specificity was set to trypsin with maximally 2 missed cleavages allowed. Minimum peptide length was set to 7, and the maximum peptide mass was set to 4600 Da. To determine higher missed cleavage rates, a search using a missed cleavage rate of 5 was used for the trypsin amount optimisation experiment. To ensure high confidence identifications, peptide-spectral matches, peptides, and proteins were filtered at a less than 1% false discovery rate (FDR). The ‘match between runs’ feature was off. Default settings were applied to the remaining parameters.

The partial-labelling rate was calculated as the ratio between the number of PSMs which had been labelled but had one or more unlabelled sites, and the total number of fully labelled PSMs. The overlabelling rate was calculated as the ratio between the number of PSMs, which had one or more labelled serine/threonine/histidine residues, and the total number of fully labelled PSMs.

DDA-TMTpro data were searched using Fragpipe^35^ (v 20.0) using a modified “TMT18-Phospho” workflow. Raw data were searched against the human proteome sequence database retrieved from SwissProt on November 6, 2020, decoy sequences and contamination sequences were automatically added by Fragpipe. TMTpro (H), Phosphorylation (S, T, Y), Oxidation (M) and Acetylation (protein N-term) were set as variable modifications, and TMTpro (K), TMTpro (peptide N-term) were set as fixed modifications. Enzyme specificity was set to trypsin with maximally 2 missed cleavages allowed. Minimum peptide length was set to 7, and the maximum peptide mass was set to 5000 Da. MSBooster^36^ rescoring, Percolator^37^ for PSM validation, PTMProphet^38^ for modification localisation and TMT-Integrator with Philosopher^39^ for reporter ion quantification were enabled and MSstats files were generated.

Mass spectrometry data have been deposited to the ProteomeXchange Consortium via the PRIDE^40^ partner repository with the dataset identifier PXD056902 (username, password: qjfWSQVIhmzE).

### Bioinformatic Analysis

Phosphosite quantification and differential expression analysis were performed using MSstatsPTM^41^, with normalisation and MBimputation enabled. CV calculations were performed using “proteomicsCV” package with the raw outputs from the Fragpipe search and non-phosphorylated peptides were omitted from the analysis^42^. Correlation coefficients were calculated using the Spearman’s rank method, with missing values replaced by 0. The clustering method to plot the heatmap was “ward.D2” without scaling, missing values were removed before analysis. Principle Component Analysis was calculated using the “prcomp” function in R. Phosphosites with missing values were removed before analysis. Kinase activity enrichment was performed using PTM-SEA^27^, with the iKiP database and the ranking metric - log10(Adj.P -value)*Sign(log2FC) calculated with the pairwise comparison results from MSstatsPTM^43^. The output from the PTM-SEA analysis was filtered to select kinases with a significant (adjusted p-value < 0.05) normalised enrichment score (NES) in at least one comparison for the low plex analysis. Using these data, kinase activity rankings were generated and presented as a radar plot.

### Statistics and Reproducibility

Optimisation experiments were performed in triplicate except for the lysis buffer inhibitor experiment (n=2) due to significant detrimental effects on the overall system performance, and the optimisation of sample input for multiplexed phosphopeptide enrichment (n=1).

Linear regression and unpaired two-sided t-test were performed in Graphpad Prism (GraphPad Software, USA). T-test results are presented in the figures as follows: ns: P > 0.05, *: P ≤ 0.05, **: P ≤ 0.01, ***: P ≤ 0.001, ****: P ≤ 0.0001.

## Data availability

Mass spectrometry data have been deposited to the ProteomeXchange Consortium via the PRIDE^40^ partner repository with the dataset identifier PXD056902.

## Acknowledgements

The UCL Cancer Institute Translational Technology Platforms (TTPs) were supported by the Cancer Research UK (CRUK) UCL City of London Centre Award (CTRQQR-2021\100004C416/A25145). We would like to acknowledge the support of the Proteomics Research TTP and the Flow Cytometry TTP. Personal funding to E.J.G was from the CRUK Non-Clinical Training Award (A31279). The patient-derived GSC lines (GCGR-E37, GCGR-E53, GCGR-L12 and GCGR-L21) were made available through the Glioma Cellular Genetics Resource funded by CRUK Accelerator Award (A21992).

## Author Contributions

E.J.G. and X.C. designed and performed experiments, analysed the data and wrote the manuscript. A.B. maintained instrumentation and provided scientific input. S.S. provided initial conceptualisation, designed experiments, supervised the study and wrote the manuscript.

## Supplementary Figures

**Supplementary Fig. 1.**
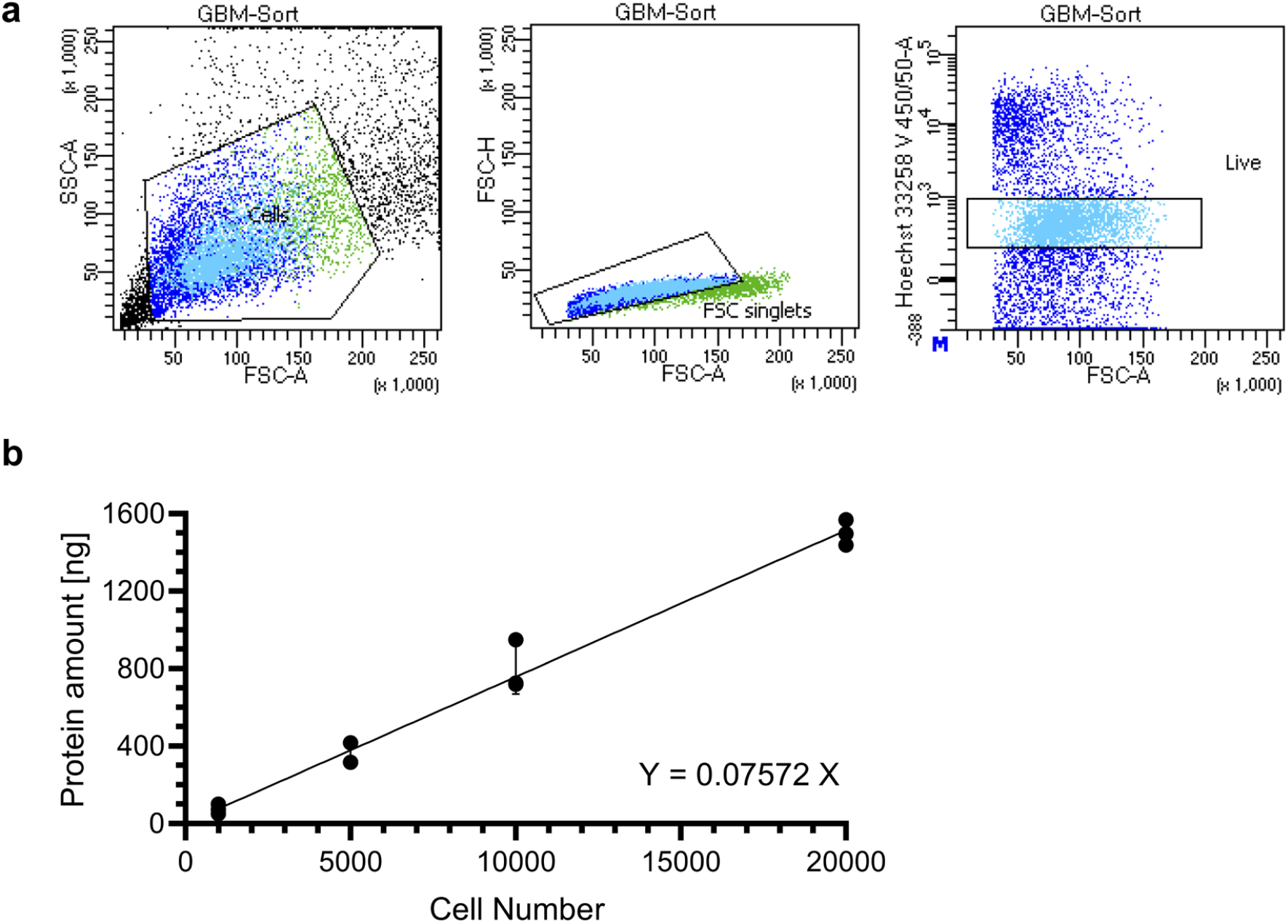
Quantification of protein yield from 1,000 GBM-E37 cells. **a** A representative FACS gating strategy to isolate live single cells. **b** Protein concentration curve for the calculation of protein amount in the GBM-E37 cell line. Data were calculated from a Micro BCA experiment using an independent standard curve and linear regression through (0, 0). A sample of 1,000 cells contains approximately 75 ng of protein.

**Supplementary Fig. 2.**
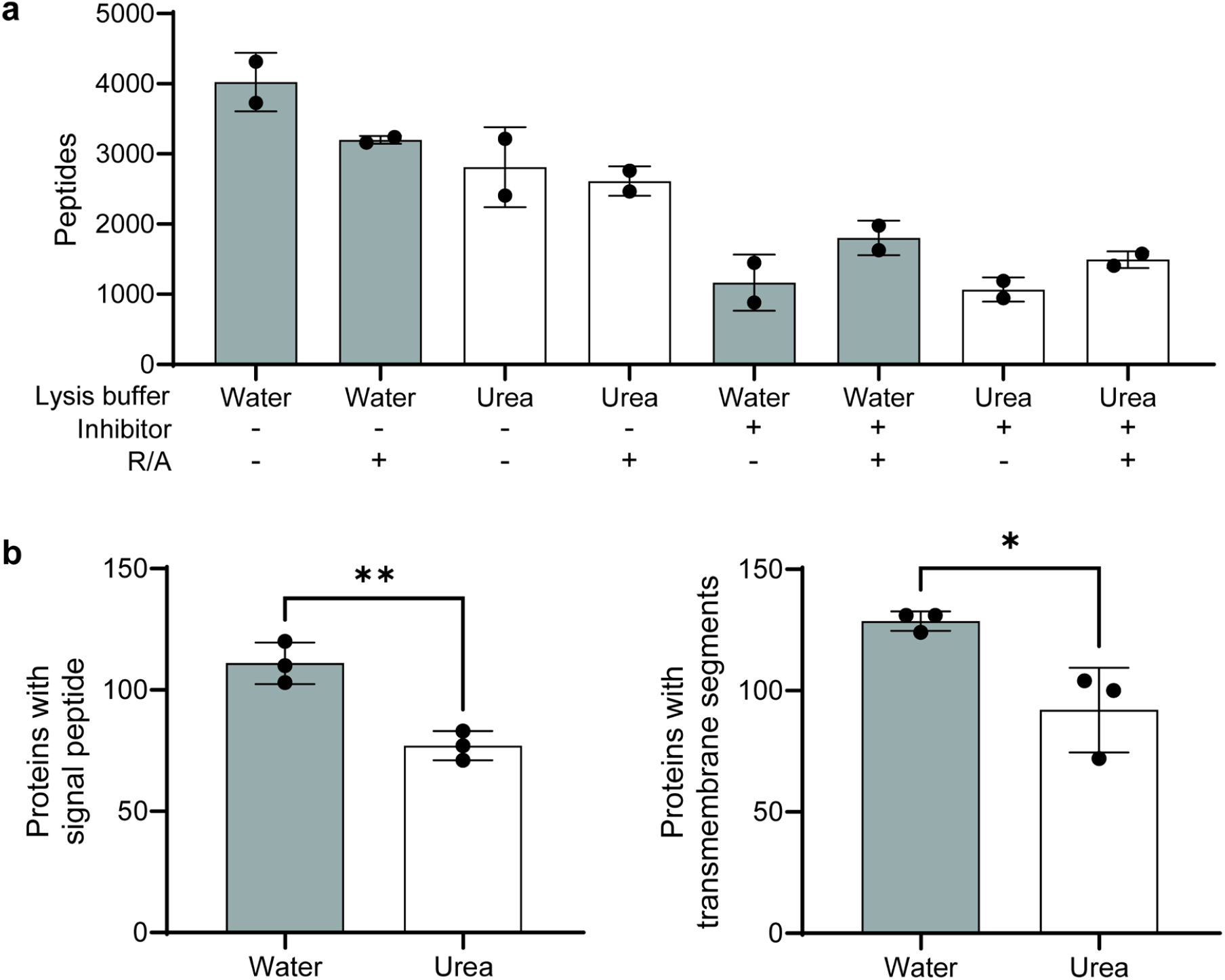
Optimisation of low-input sample lysis. **a** Evaluation of the effect of protease inhibitors in combination with reduction and alkylation (R/A) on digestion efficiency. **b** The number of identified proteins with a signal peptide (left) and transmembrane segment (right) with different lysis methods.

**Supplementary Fig. 3.**
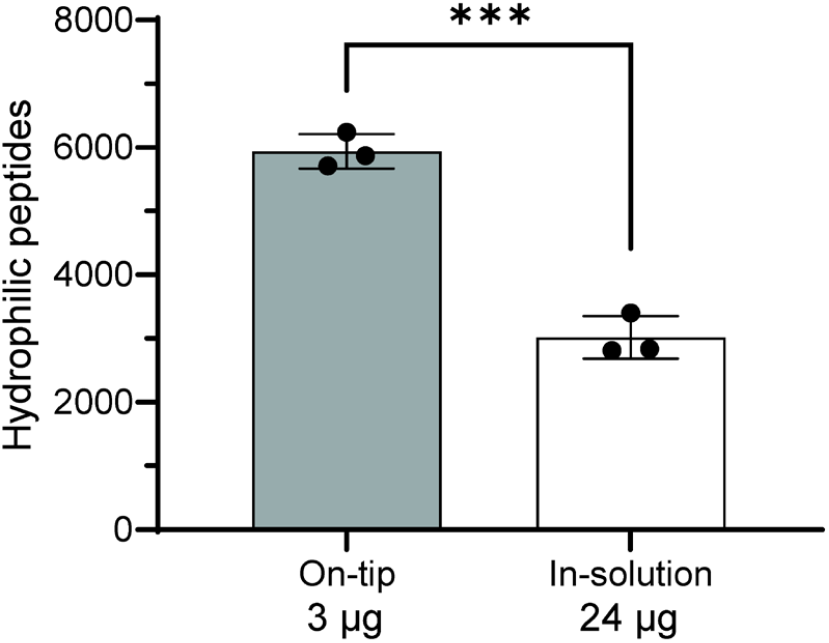
Comparison of on-tip and in-solution labelling methods. The number of hydrophilic peptides identified with the on-tip and the in-solution method.

**Supplementary Fig. 4.**
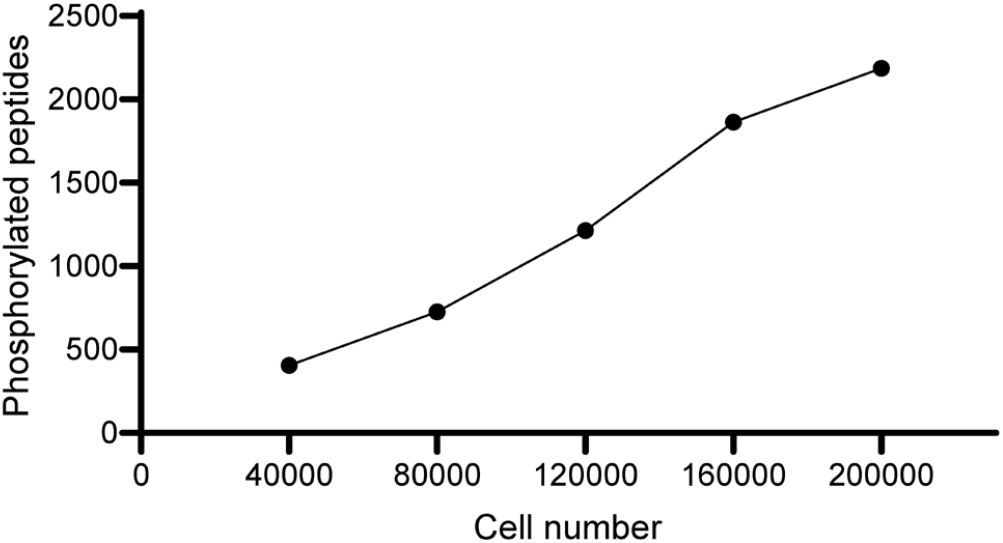
Optimising sample amount for multiplexed phosphopeptide enrichment. The relationship between different cell number input and identified phosphorylated peptides.

**Supplementary Fig. 5.**
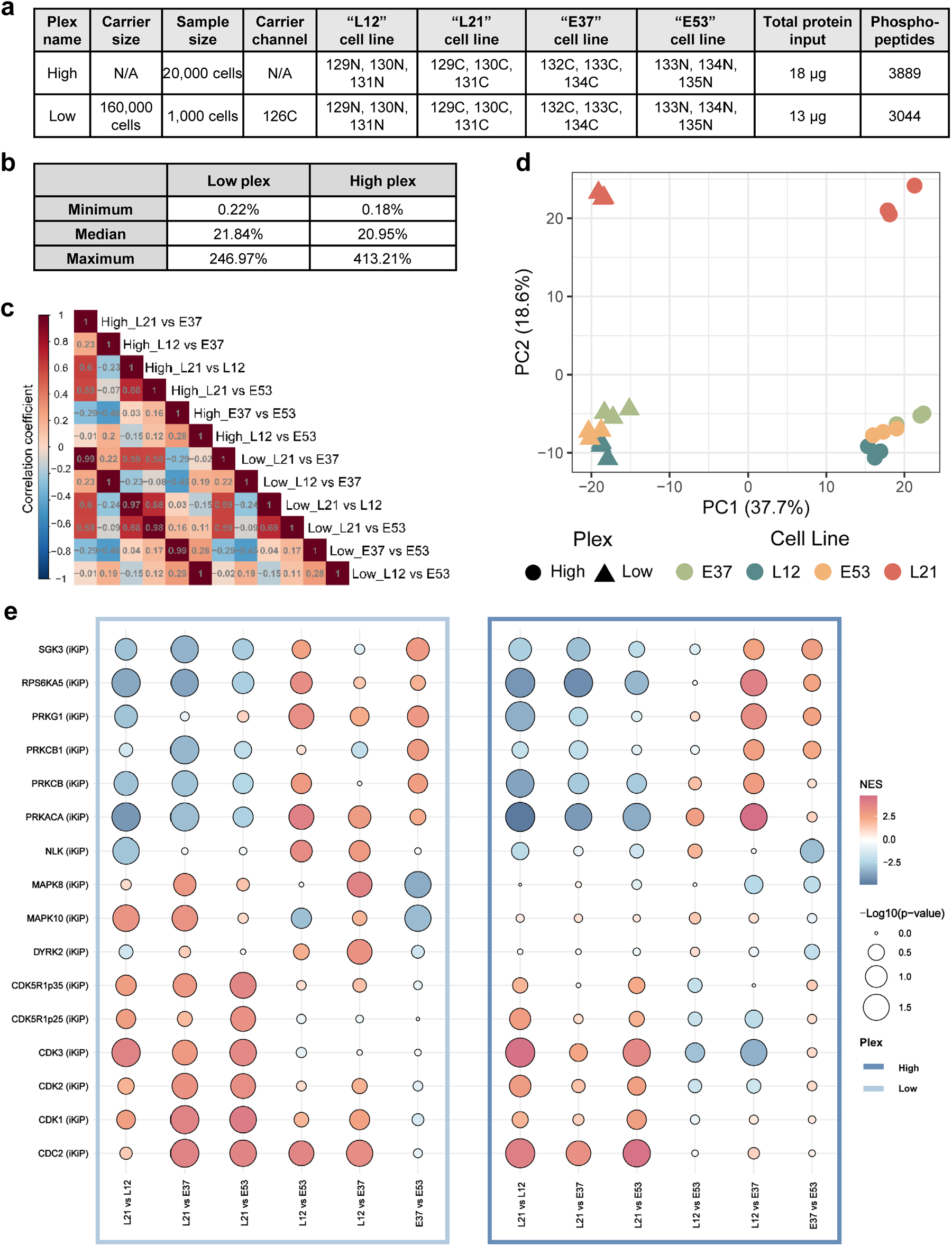
Comparison of the SPARCE workflow using different input amounts. **a** The experimental design to investigate the differences between four patient-derived glioblastoma cell lines. **b** The maximum, minimum, and median CV values of the quantified phosphosites from both plexes. **c** The correlation heatmap between comparison groups within the two plexes. **d** The principal component analysis (PCA) results based on phosphosite abundance, x-axis is PC1 and the y-axis is PC2. **e** The kinase activity profiles of different cell lines, inferred and ranked using PTM-SEA.

